# Improved interpretability in LFADS models using a learned, context-dependent per-trial bias

**DOI:** 10.1101/2025.10.03.680303

**Authors:** Nishal P. Shah, Benyamin Abramovich Krasa, Erin Kunz, Nick Hahn, Foram Kamdar, Donald Avansino, Leigh R. Hochberg, Jaimie M. Henderson, David Sussillo

## Abstract

The computation-through-dynamics perspective argues that biological neural circuits process information via the continuous evolution of their internal states. Inspired by this perspective, Latent Factor Activity using Dynamical systems (LFADS, [1]) identifies a generative model consistent with the neural activity recordings. LFADS models neural dynamics with a recurrent neural network (RNN) generator, which results in excellent fit to the data. However, it has been difficult to understand the dynamics of the LFADS generator. In this work, we show that this poor interpretability arises in part because the generator implements complex, multi-stable dynamics.

We introduce a simple modification to LFADS that ameliorates issues with interpretability by providing an inferred per-trial bias (modeled as a constant input) to the RNN generator, enabling it to contextually adapt a simpler dynamical system to individual trials. In both simulated neural recordings from pendulum oscillations and real recordings during arm movements in nonhuman primates, we observed that the standard LFADS learned complex, multi-stable dynamics, whereas the modified LFADS learned easier-to-understand contextual dynamics. This enabled direct analysis of the generator, which reproduced at a single-trial level previous results shown only through more complex analyses at the trial average. Finally, we applied the per-trial inferred bias LFADS model to human intracortical brain computer interface recordings during attempted finger movements and speech. We show that modifying neural dynamics using linear operations of the per-trial bias addresses non-stationarity and identifies the extent of behavioral variability, problems known to plague BCI. We call our modification to LFADS as “contextual LFADS”.

## 1 Introduction

The dynamical systems perspective is increasingly being adopted to understand how neuronal circuits in the brain represent and compute information [2–4]. This perspective states that neural activity embeds the evolution of a latent state, which starts from an initial condition and evolves according to recurrent dynamics of the neural circuit as well as inputs from other brain regions. A complete characterization of neuronal activity under this framework entails estimating the dynamical system, as well as the initial conditions and input that drive it. While the latent states give an interpretable description of the high-dimensional neural activity, the components of the dynamical system provide mechanistic interpretation of the neural circuit.

When it comes to estimating such a system, linear dynamical systems provide a simple and interpretable model in terms of the separable modes that evolve independently. While easy to understand, the linear models may not be the best fit to neural data. Subsequently, variants of the linear model with discrete switching [5] or smooth interpolations [6, 7] improve model fit while still being interpretable. However, these approaches impose assumptions about how dynamics vary across task conditions, embedding those assumptions directly into the model architecture.

Alternatively, flexible models such as RNNs [1, 8, 9] or transformers [10–14] do not make such assumptions and achieve state-of-the-art accuracy in fitting to neural data. A prominent example of this is LFADS [1], an unsupervised sequential auto-encoder for neural time series. LFADS uses a recurrent neural network (RNN) generator to model the neural population dynamics, extracting a set of latent factors that summarize high-dimensional neural spiking on single trials. By making an explicit dynamical systems assumption, LFADS infers smooth latent trajectories that capture the neural computations within each trial, leading to improved single-trial representations for downstream decoding or analysis. In practice, LFADS models have shown state-of-the-art performance in denoising neural data and inferring latent firing rates, providing researchers with a window into the “neural manifold” underlying complex behaviors [1, 15, 16].

Interpreting the dynamics learned by an RNN typically requires a reverse engineering step in which one identifies fixed points and analyzes the linearized dynamics around them [17, 18]. While this approach can in principle provide mechanistic insights [19], it has yielded limited success in practice for LFADS thanks to LFADS generators often developing complex, *multi-stable dynamics*. This in turn makes fixed-point finding and linearization far less useful.

Here we propose a straight-forward modification that gives the LFADS generator access to an inferred low-dimensional per-trial bias, modeled as an additional constant input. This bias parameterizes a (low-dimensional) family of RNNs and thereby subtly adapts the RNN dynamics on a per-trial basis, resulting in *contextual dynamics* that are often simpler to interpret than the complex, often multi-stable dynamics of the original LFADS architecture. Further, the per-trial bias gives a low-dimensional embedding of the family of dynamical systems (parameterized by each bias), which can be used for additional analyzes as well as for better BCI decoder estimation or improved neural data analysis. The main contributions of our work are:

1. We identify complex, often multi-stable dynamics as a fundamental barrier in using LFADS to understand dynamics in neural recordings.
2. We introduce a new modification of LFADS with a per-trial bias for learning contextual dynamics, which are often easier to interpret. The modification is straightforward and can be easily incorporated into existing implementations [20–22].
3. In recordings from monkey motor cortex during reaching, we show that our model recovers known structure in the neural dynamics and reveals novel, condition-dependent changes in oscillation frequency at the single-trial level [23].
4. We study the per-trial bias embedding space and show that simple geometrical operations of per-trial bias can mitigate the effects of non-stationarity and identify the extent of behavioral variability in human intracortical BCI recordings.

## 2 Related works

Task RNNs [24–26] are artificial recurrent networks trained to solve cognitive tasks in simulation, and are useful to build hypotheses about how the brain can solve these tasks. Task conditions (such as stimulus, task type or task period) are often provided to the RNNs using constant inputs, similar to the per-trial bias in our model. Similar to our analysis, Driscoll et al., 2024 [24] perform interpolation operation between the constant inputs (bias) for two different conditions and study how the dynamics change smoothly with the interpolation.

Several approaches have sought to improve the interpretability of RNNs fit to neural recordings by introducing structural constraints. One popular strategy constrains the recurrent dynamics to evolve within a low-dimensional subspace, enabling analytical insights into their behavior [27–31]. However, such low-rank constraints may limit biological realism. Other work has leveraged RNNs with piecewise-linear activations [32–37], which allow tractable analysis of fixed points and cycles. Neural ODE models, which learn continuous-time dynamics by predicting state derivatives rather than next states, have also been shown to enhance interpretability [38, 39]. In contrast to these structural modifications, our approach retains standard nonlinear RNN modules such as GRUs [40], and instead improves interpretability by constraining the per-trial contextual bias to vary along a low-dimensional manifold. This approach assumes that while the neural activity within each trial may be high-dimensional, the across-condition variability is captured compactly in a lower-dimensional space.

Finally, Jacobian Switching Linear Dynamical System (JSLDS, [9]) improves interpretability by co-training the LFADS RNN with a switching linear dynamical system. At each time point, this model identifies a single fixed point for linearizing the dynamics and this fixed point can change over the course of a trial. In contrast, our approach can learn more complex structure (multiple fixed points) that remains fixed for the course of the trial.

## 3 Methods

We begin by reviewing the standard LFADS model and then introduce our proposed modification incorporating a per-trial contextual bias (Figure 1).

**Figure 1:**
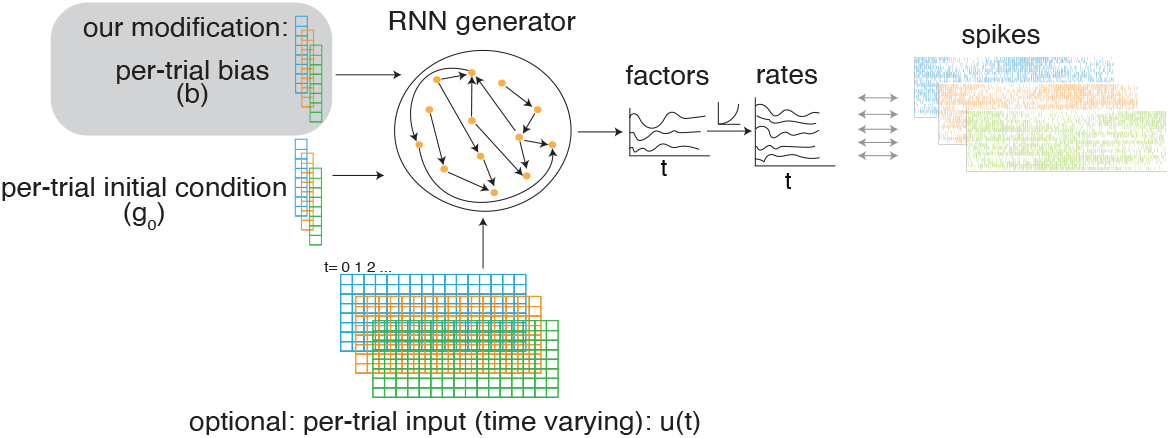
The contextual LFADS model. Recorded spikes are modeled as samples from a Poisson firing rate. Log-firing rate is linearly read out from the RNN state with an optional bottleneck of low-dimensional factors. In the standard LFADS, the RNN starts from an initial condition and optionally driven by a low-dimensional time-varying input (both inferred from data from encoders, not shown). Our modification of LFADS provides an additional inferred per-trial bias (gray box) which is modeled as a constant input to the RNN.

### 3.1 The Standard Latent Factor Analysis via Dynamical Systems (LFADS) Model

LFADS [1] is a sequential variational autoencoder designed to infer smooth latent dynamics from noisy, high-dimensional neural spike recordings. The model represents each trial’s neural activity as the output of a nonlinear dynamical system, simulated by an RNN. For trial number *i*, the RNN generates latent states **g**_*i,t*_ from an initial condition **g**_*i*,0_ and optional inferred inputs **u**_*i,t*_ (both are outputs of separate probabilistic encoders, not shown).

For trial *i* with observed spike counts **x**_*i*,1:*T*_, the generative process is:

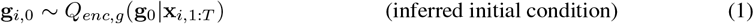

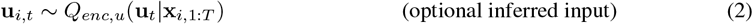

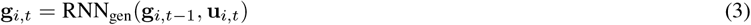

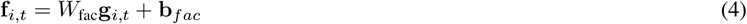

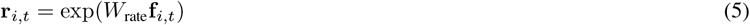

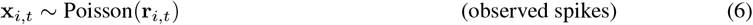

where the optional time-varying factors **f**_*i,t*_ provide a low-dimensional representation of population activity, and **r**_*i,t*_ are the Poisson firing rates. The encoders (*Q*_*enc,g*_, *Q*_*enc,u*_) are bidirectional RNNs followed by linear mapping to a lower dimension. We use reparametrization trick to generate samples for initial condition and time-varying input. For example, the initial condition encoder predicts mean (**g**_*i*,0,*µ*_) and standard-deviation (**g**_*i*,0,*σ*_) and samples are generated by **g**_*i*,0_ = **g**_*i*,0,*µ*_ + *ϵ***g**_*i*,0,*σ*_ with *ϵ* distributed as standard Gaussian. See Pandarinath et al. 2018 [1] for further details.

### 3.2 Modification of LFADS with a Per-Trial Bias

To improve interpretability and reduce model complexity, we introduce a per-trial contextual bias **b**_*i*_. The per-trial bias is modeled as an constant input to the RNN, in addition to the optional time-varying inferred inputs **u**_*i,t*_. The RNN update equations are given by:

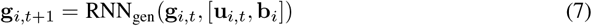

where the input and bias are concatenated ([**u**_*i,t*_, **b**_*i*_]) at each time-step. Note that the per-trial bias is inferred by an encoder from the observed spike train:

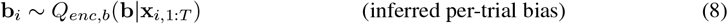

Similar to the encoder for initial condition, the bias-encoder (*Q*_*enc,b*_) is a bidirectional RNN, and reparameterization trick is used to generate samples. The per-trial bias is typically lower dimensional (dimensionality ≤10) than the RNN hidden state dimensions. The training objective is presented in Appendix section 6.1, details on model fitting is presented in Appendix section 6.2 and hyperparameters are listed in Appendix section 6.4.

The per-trial bias allows the generator to implement trial-specific dynamics via the (potentially smooth) modulation of the dynamical landscape. Although the mean of the time-varying input could, in principle, capture similar information as the per-trial bias, the full complexity of a time-varying inferred input is often unnecessary for many analyses. Moreover, commonly used priors on the input—such as Gaussian or smooth autoregressive priors—can obscure the mean, making it more difficult to learn explicitly. Empirically, the result is often more interpretable using an inferred bias, where each **b**_*i*_ moves a single fixed point rather than relying on multi-stable autonomous dynamics or time-varying dynamics resulting from time-varying inputs.

## 4 Results

### 4.1 Interpretable Dynamics in a Simulated Pendulum

To evaluate the interpretability of the proposed method, we simulated a system with known ground truth dynamics: a family of damped pendulums (Figure 2A). This example is constructed to make the point that while a single pendulum should result in simple dynamics around a single fixed point, a family of pendulums will result in multi-stable dynamics. When modeling neural data with LFADS we will not know which scenario we are in.

**Figure 2:**
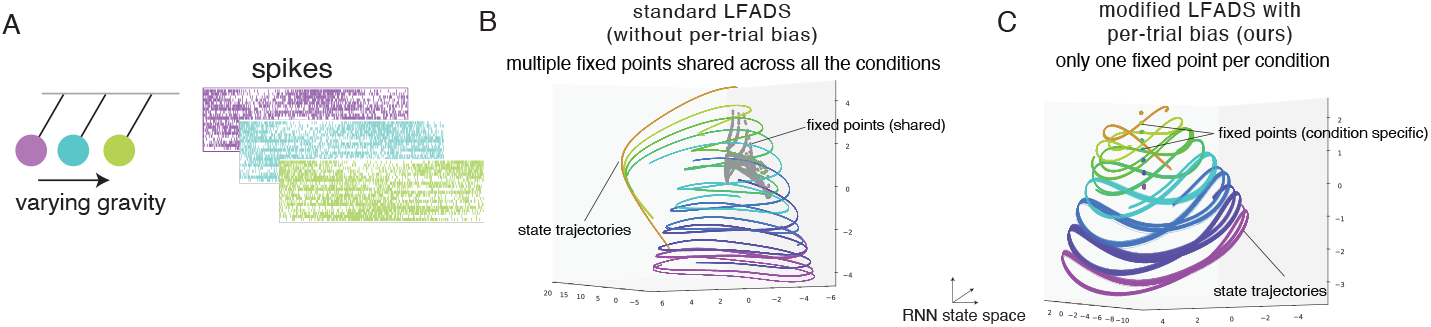
Simulated pendulam. (A) Task conditions correspond to distinct values of gravity, which changes the oscillation frequency. Neural activity embeds oscillations of the pendulum. Multiple trials generated for each gravity condition. (B) Standard LFADS. Colored lines indicate RNN hidden state trajectories (projected to top three principal components), colors indicate oscillation frequency/gravity. Gray dots indicate fixed points, which are shared across all conditions. (C) Same as (B) for contextual LFADS with inferred per-trial bias. The dots indicate fixed points, and for a given task condition only the fixed points with the corresponding color are present.

The pendulum’s state was represented as 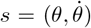, where *θ* denotes the angle. Starting from an initial state *s*_0_ = (*θ*_0_, 0), the state evolved according to the nonlinear differential equation given by 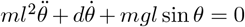, with *m* = 1, *l* = 5 and *d* = 1. The activity of 192 simulated neurons (*n*) embedded the state of the pendulum using a mapping *n* ~Poisson(*max*(*Ks* + *µ*, 0)). We simulated a family of pendulums under 8 distinct gravity conditions by changing the values of *g* from 0.5 to 4.5 in steps of 0.5. For each condition, we generated 125 trials of 300 sample duration, with 75 used for training, 25 for validation and 25 for testing.

We trained both the standard LFADS model and the contextual LFADS model incorporating per-trial biases on the simulated dataset. As the ground-truth dynamics are autonomous, we did not have the optional time-varying input for both models. We observed that the both models fit the data well and showed oscillations in the hidden state of the generator (Figure 2B,C).

To understand the learned dynamics, we “reversed engineered” the learned RNN to identify approximate fixed points of the state update function [17, 18]. Specifically, the fixed points were dentified as neural states where **g**\* ≈ RNN_gen_(**g**\*, **u**_*i,t*_ = 0) for the standard LFADS and **g**\* ≈ RNN_gen_(**g**\*, [**u**_*i,t*_ = 0, **b**_*i*_]) for the LFADS model with per-trial bias. Details of fixed point finding are presented in Appendix (Section 6.3).

We observed that the standard LFADS model implemented a cluster of fixed points which were shared across all conditions (Figure 2B, gray), with each fixed point associated with dynamics of a distinct oscillatory frequency. Consequently, the initial condition of the RNN not only determined the initial state of the pendulum, but also had to implicitly selected the correct pendulum from the family of pendulums. To accomplish this, the RNN implemented a **multi-stable dynamical** landscape.

In contrast, we observed that the LFADS model with per-trial biases implemented a single fixed point per trial. Within each condition, both the inferred bias and the corresponding fixed-point location remained consistent across trials (Figure 2C). As the gravity level varied, the fixed point location and bias moved smoothly along a one-dimensional manifold, effectively recovering the underlying scalar parameter. This structure is described as **contextual dynamics**, in which the RNN dynamics are explicitly modulated by inferred contextual inputs, leading to an interpretable representation of condition-specific variability. In the next section, we extend these observations to real data.

### 4.2 Neural dynamics during reaching task in monkeys

We further evaluated contextual LFADS on the “Maze dataset” [23], in which a monkey performed straight and curved reaches while sitting in front of a computer screen (Figure 3A,B). The dataset consists of 107 reach conditions, with 14 *−* 39 trials per condition (average 18.3 trials per condition). Each trial spans 180 ms before to 450 ms seconds after the go cue.

**Figure 3:**
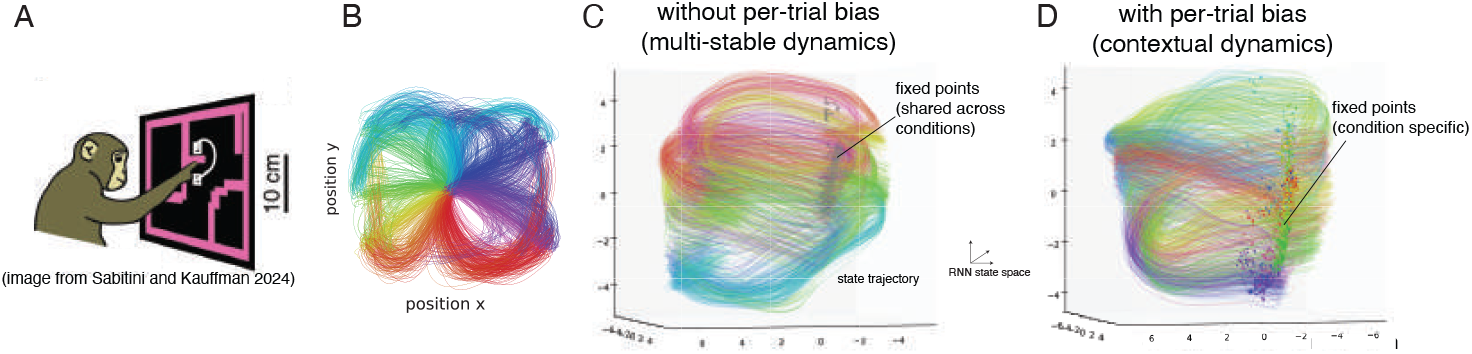
Maze task. (A) Task setup showing monkey reaches around obstacles to reach targets, resulting in straight or curved reaches. Figure from [41]. (B) 2D arm trajectories. Colors indicate initial reach direction. (C) RNN hidden state visualization (along top 3 PCs) for the standard LFADS model. Lines indicate state trajectory, colors indicate initial reach direction (same as (B)). Gray dots indicate the shared fixed points across all conditions. (D) Same as (C), for contextual LFADS with inferred per-trial bias. Colored dots indicate fixed point locations. For a given trial, only a single fixed point was active (shown in corresponding color).

As with the pendulum simulation, we observed that both standard and contextual LFADS models identified similar latent dynamics and achieved comparable kinematic reconstruction accuracy (linear regression, Standard LFADS: 0.842, SD: 0.008 vs Contextual LFADS: 0.844, SD: 0.012). The standard LFADS model exhibited multi-stable dynamics, characterized by a shared cluster of fixed points across conditions (Figure 3C). In contrast, the modified LFADS model with per-trial bias exhibited contextual dynamics, with one fixed point per condition, leading to a more interpretable organization of trial-to-trial variability (Figure 3D).

A key scientific question that can be addressed with this dataset is which aspects of neural dynamics are invariant across reach conditions and which are condition-dependent. Churchland and Shenoy, 2012 [23] argued that starting from a reach-specific preparatory activity (corresponding to initial condition), neural activity follows shared condition-invariant dynamics. Recently, Sabatini and Kauffman, 2024 [41] showed that all properties of the dynamics except rotation frequency are condition-specific. We revisit this question using the contextual LFADS model.

We observe that the inferred per-trial biases captured the circular geometry of reach directions, with biases corresponding to similar directions located nearby (Figure 4A). As the bias varied smoothly around the circle, the associated fixed point locations varied smoothly in the RNN state space (Figure 4B).

**Figure 4:**
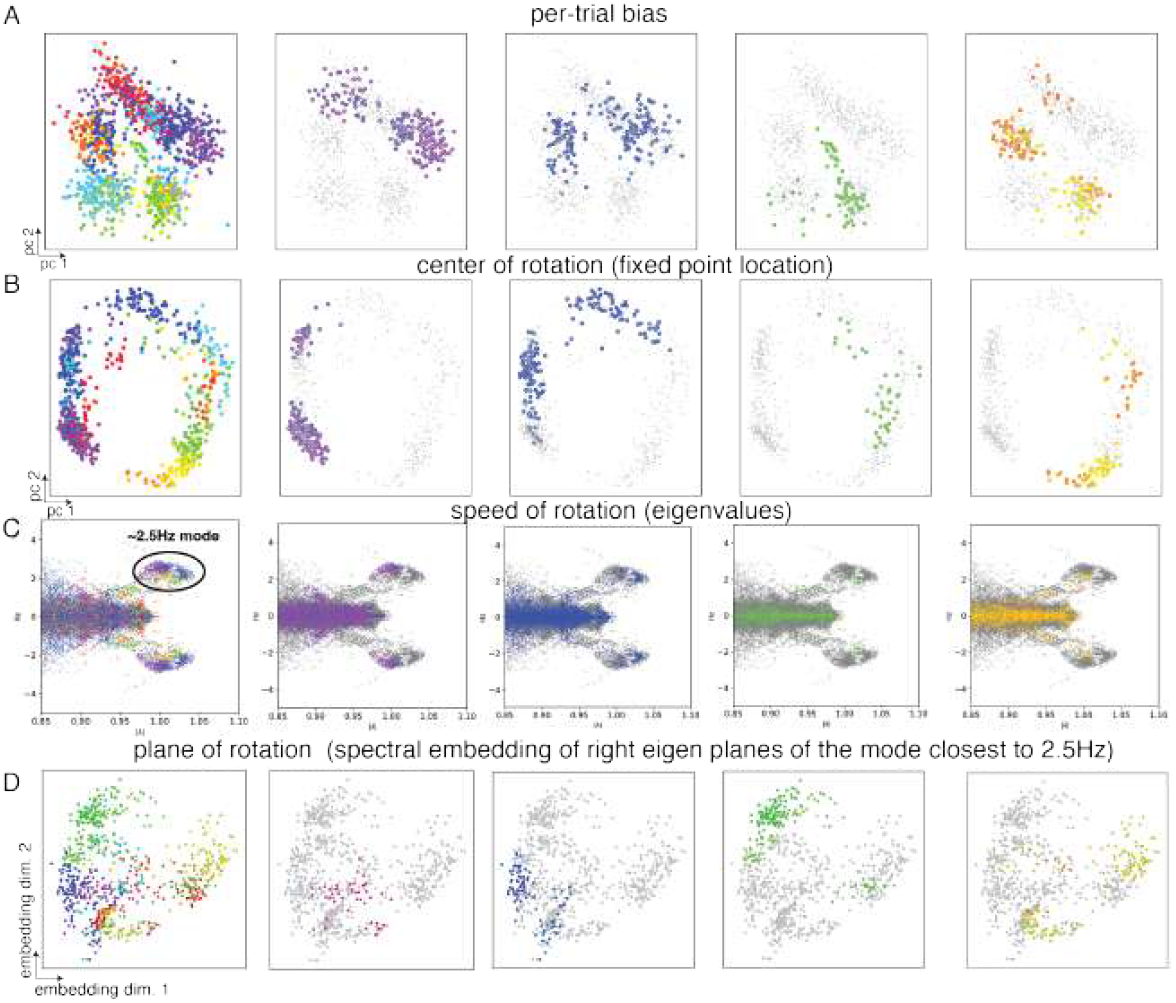
Contextual dynamics in the Maze task. Reach condition–dependent variation in different components of the dynamics system. Each dot represents a single trial. Color indicates the initial reach direction across all trials. In the first column, reach direction of all trials is indicated. In the subsequent columns, only trials within with a narrow range of reach angles is indicated (by color), while other trials are in gray. (A) Per-trial bias, projected onto its top two principal components. (B) Fixed-point location, projected onto its top two principal components. (C) Oscillation frequency, visualized via the real and imaginary parts of the eigenvalues across trials. (D) Two-dimensional spectral embedding of planes of rotation for the dominant oscillatory mode near 2.5 Hz. Similarity between trials was quantified based on how well the RNN state trajectory of one trial could be reconstructed using the eigen-planes from another. Similarity of center of rotation (B) and plane of rotation (D) for similar reach directions is consistent with [41]. The condition-specific changes in eigenvalues (C) has not been reported before.

Next, we studied the linearized dynamics around the fixed points, approximating *RNN*_gen_(**x**, [**u**_*i,t*_ = 0, **b**_*i*_]) ≈ *A*(**b**_*i*_)(**x** − **x**_fixed point_(**b**_*i*_)) + **x**_fixed point_(**b**_*i*_). Across different per-trial biases, we compared eigenvectors and eigenvalues of *A*. The real and imaginary components of the eigenvalue indicate the decay and frequency of rotations, and the eigen-planes (span of a pair of conjugate eigenvectors) indicate the planes in which these rotations lie.

Across different trials, there was a consistent 2.5 Hz oscillation mode in the eigenvalues (see the cluster in Figure 4C). The plane spanned by eigenvectors with eigenvalue closest to the 2.5 Hz frequency varied continuously with reach direction (Figure 4D). A modest but consistent modulation of the eigenvalue (oscillation frequency) near the 2.5 Hz mode was also detected across conditions (Figure 4C). While condition-dependent changes in fixed-point locations and eigen-planes have been previously reported [41], the eigenvalue modulations were not captured by earlier analyses relying on trial averaged data. Our single-trial analysis reveals a more nuanced view: all dynamical properties exhibit condition-specific variation, though the degree of variability differs across them (Table 1). We emphasize that contextual LFADS aided us in arriving at these results through no specific knowledge or hypotheses concerning these data (beyond trial structure).

**Table 1:**
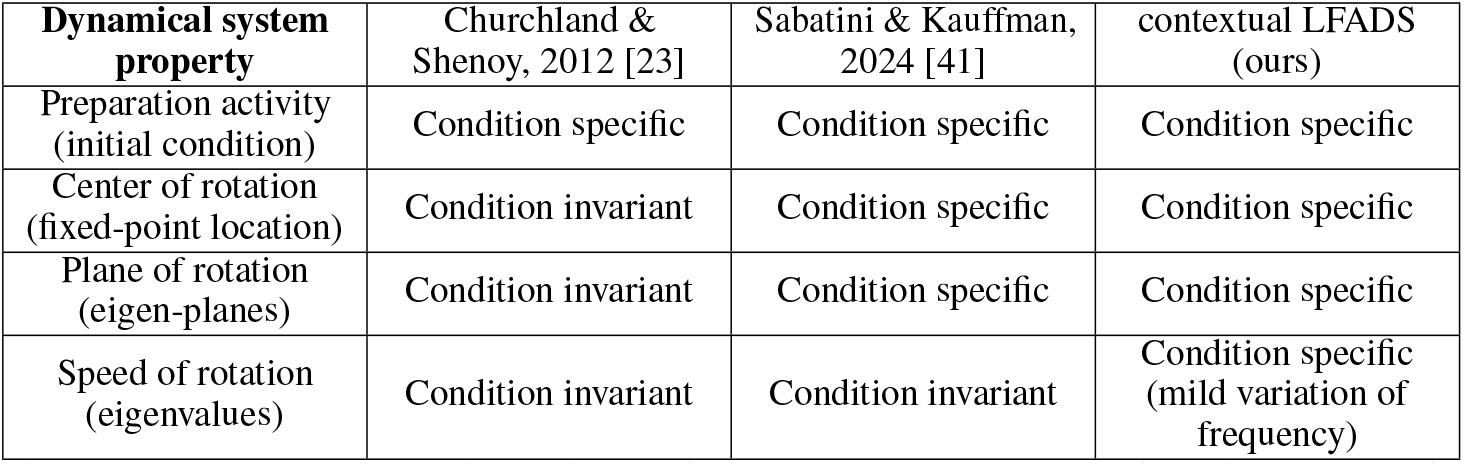
Dynamical systems understanding of the neural activity in Maze reaching dataset. Note that contextual LFADS uses single-trial analysis whereas the other two use trial-averaged analysis.

| Dynamical system property | Churchland & Shenoy, 2012 [23] | Sabatini & Kauffman, 2024 [41] | contextual LFADS (ours) |
| --- | --- | --- | --- |
| Preparation activity (initial condition) | Condition specific | Condition specific | Condition specific |
| Center of rotation (fixed-point location) | Condition invariant | Condition specific | Condition specific |
| Plane of rotation (eigen-planes) | Condition invariant | Condition specific | Condition specific |
| Speed of rotation (eigenvalues) | Condition invariant | Condition invariant | Condition specific (mild variation of frequency) |

### 4.3 Data augmentation using contextual LFADS makes iBCI decoders robust to non-stationarities

A major challenge in intracortical brain-computer interface (BCI) studies is the issue of non-stationarity, where neural recordings corresponding to the same intended movements drift over time. To investigate whether contextual LFADS could mitigate the effects of non-stationarity, we analyzed data from a participant ‘T5’ (enrolled in the BrainGate2 clinical trial) with two 96-channel Utah arrays surgically placed in the hand-knob region of the precentral gyrus. During the experiment, the participant attempted combinatorial movements of two finger groups: the thumb (one degree of freedom) and all other fingers (second degree of freedom). Each group was independently cued to flex, extend, or remain at rest. Trials alternated between transitions from rest to a cued gesture and back to rest. A total of 966 trials were collected, 52-54 trials for each of the 16 movements, and 107 trials without movement.

We focus on the goal of learning a linear finger position (continuous valued) decoder for the final quarter of the trials collected during a recording session. First, we evaluated the performance of a decoder trained on Gaussian-smoothed neural activity. Using the trials only from the final quarter yielded significantly lower accuracy compared to using all trials (*R*^2^ of 0.24 vs. 0.46, Table 2), likely due to reduction in training data despite of an improved match in data distribution.

**Table 2:**
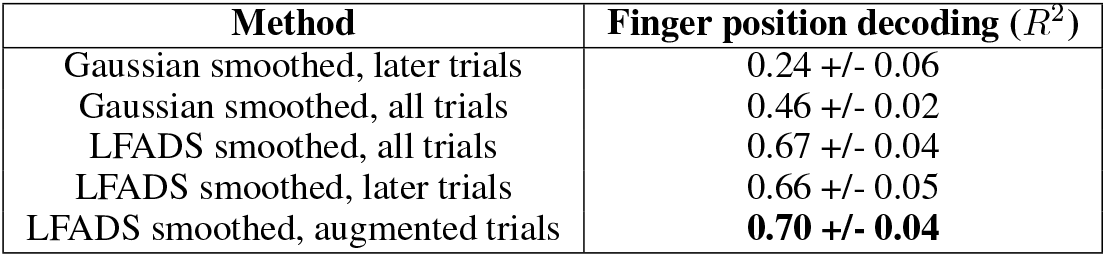
Using contextual LFADS for data augmentation improves decoder accuracy in the presence of non-stationarities. Performance of “LFADS smoothed, augmented trials” is significantly better than “LFADS smoothed, later trials” (paired t-test, *p* < 0.01).

| Method | Finger position decoding ($R^2$) |
| --- | --- |
| Gaussian smoothed, later trials | 0.24 +/- 0.06 |
| Gaussian smoothed, all trials | 0.46 +/- 0.02 |
| LFADS smoothed, all trials | 0.67 +/- 0.04 |
| LFADS smoothed, later trials | 0.66 +/- 0.05 |
| LFADS smoothed, augmented trials | **0.70 +/- 0.04** |

Training and testing the decoder on LFADS-inferred firing rates improved prediction accuracy, as expected from prior work [1]. Moreover, the discrepancy between training exclusively on the final quarter versus all trials was reduced (0.67 vs. 0.66, Table 2), suggesting that improved smoothing yields decoders that are robust to non-stationarities.

Finally, we used contextual LFADS model to transform data collected early in the session to later in the session. Analysis of the top two principal components of the inferred biases revealed a drift over the course of the session (Figure 5C). To compensate for this drift, per-trial biases and initial condition from early-session trials were systematically shifted to match the corresponding parameters for later-session trials.

**Figure 5:**
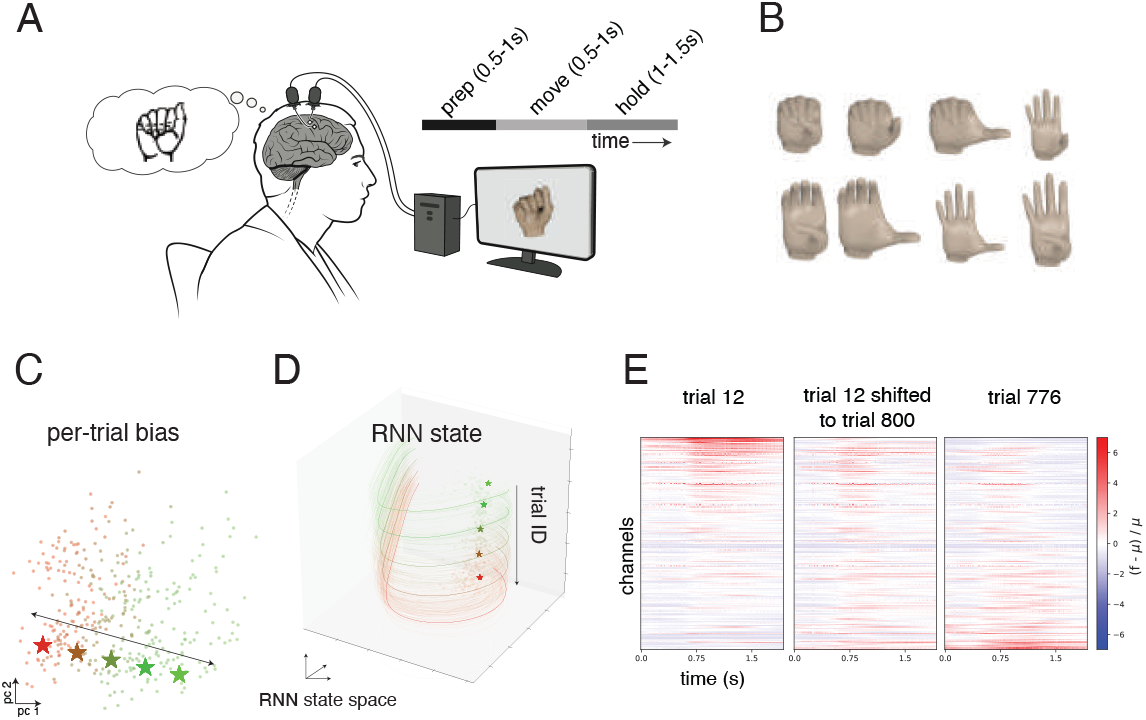
Using contextual LFADS for data augmentation improves decoder robustness to non-stationarities. (A) Intracortical BCI participant attempted finger movements as neural activity was recorded using Utah arrays. (B) The corpus of finger movement targets, with two finger groups (thumb and all other fingers constrained together) cued independently to be flexed/extended/stay at rest. (C) Two dimensional PCA of per-trial bias, with trials (dots) colored by the trial number. The direction of most correlated with trial number is indicated. (D) Three dimensional PCA of RNN state. Colors indicate trial number. Moving the bias (stars in panel C) moves the RNN state and fixed point (stars in panel D and solid lines). (E) Visualizations of smoothed neural activity (relative to mean for each channel) for trial 12 (left), trial 12 shifted to trial 800 (middle) and trial 776 (trial closest to 800, with same movement as trial 12, right).

Specifically, we first applied the bias-encoder to estimate the bias for trials in training partition. Next, we estimated the average bias of training trials collected early in the session **b**_early_ and estimated the average bias of training trials collected late in the session **b**_late_. Finally, for each trial collected earlier in the session, we corrected their bias with **b**_*i*,corrected_ = **b**_*i*_ − **b**_early_ + **b**_late_. We applied a similar transformation for correcting initial conditions. No changes were applied to the time-varying inputs. Further details are provided in Appendix (Section 6.5).

The shifted biases and initial conditions were then passed through the RNN to generate drift corrected neural activity, augmenting the decoder training corpus. For a given trial, as the bias moved along the non-stationarity axis, the RNN activity and the fixed points moved along the non-stationarity axis in the RNN state space (Figure 5C, D). The augmented firing rates were then combined with non-augmented firing rates for trials later in the session to estimate the decoder.

Qualitative comparison showed that the transformed neural activity closely resembled that of a late-session trial with identical cued movement (Figure 5 E). As a result, combining LFADS-inferred rates from these transformed trials with LFADS-inferred rates from trials collected later in the session resulted in the highest decoding accuracy for the later trials (*R*^2^=0.70, Table 2).

### 4.4 Per-trial bias helps identify variability in speed of putative behavior

Another potential source of variability in neural recordings with iBCI participants is the variability in the execution speed of a given movement across trials. We tested if the contextual LFADS model (without inferred inputs) could separate trials according to movement speed. Neural activity (multi-unit threshold crossings) was recorded from two 64-channel Utah arrays implanted in the ventral precentral gyrus of an BrainGate2 iBCI participant (‘T12’) as they attempted to speak 39 phonemes, (Figure 6A) with 20 trials each (data taken from [42]).

**Figure 6:**
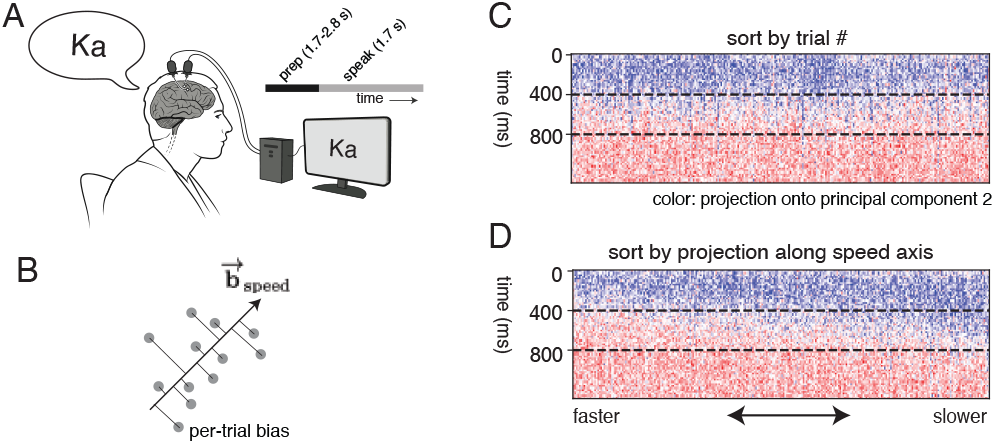
Using contextual LFADS to sort trials according to the speed of neural activity. (A) Intracortical BCI participant attempted to speak phonemes as neural activity was recorded using Utah arrays. (B) Visualization of projection of per-trial bias along a speed axis. (C, D) Visualization of first principal component of neural activity across trials, sorted by trial number (C) and projection along speed axis (D).

Projecting the neural activity along the second principal component of the condition invariant signal (i.e. neural activity over channels and time, averaged across all trials) and sorting them according to the order in which they were collected did not show any systematic structure in variation across trials (Figure 6C). Assuming that changes in the behavioral speed changes the speed of traversal of neural activity, we identified a direction in the per-trial bias space that directly affected the speed of neural trajectory traversal. For a given speed-up factor *α* (with *α <* 1 for slowdown and *α >* 1 for speedup), we identified a shift in per-trial bias space (**b**_speed,*α*_) that accelerates the neural state updates by solving the following optimization problem: **b**_speed,*α*_ = arg min_Δ**b**_ **∑** _*i,t*_[(RNN_gen_(**g**_*i,t*_, [**u**_*i,t*_ = 0, **b**_*i*_ +Δ**b**]) −**g**_*i,t*_) −*α*(RNN_gen_(**g**_*i,t*_, [**u**_*i,t*_ = 0, **b**_*i*_]) −**g**_*i,t*_)]^2^. Intuitively, we seek a shift in per-trial bias (Δ**b**) that scales the one-step RNN updates by a factor *α*.

We identified a common direction for speed 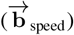 as the left singular vector of the matrix with columns corresponding to **b**_speed,*α*_ for 7 regularly spaced *α* between 0.7 and 1.3 (except *α* = 1). Sorting the trials according to their projection along this direction (i.e. sorting by 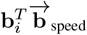) ordered them by their putative speeds in behavior, as seen from the visualization along the first principal component of the neural activity (Figure 6B,D). Hence, the contextual LFADS model helps identify the extent of behavioral variability in the neural recordings.

## 5 Discussion

We propose a modification of LFADS with an additional inferred per-trial bias provided to the RNN generator. This modification led to interpretable, contextual dynamics in the learned RNN in synthetic (Figure 2) and real monkey maze data which led to new scientific insights (Figures 3, 4). As low-dimensional bias summarizes neural dynamics, simple geometrical operations in the bias space enabled us counter non-stationarity (Figure 5) and identify the extent of behavioral variability (Figure 6)in human intracortical BCI recordings. While there may be other specialized or bespoke methods that may perform better for each of these problems, it is important to note that the method proposed in this paper is unsupervised and each of these problems is addressed in post-hoc analyses.

A key empirical observation was that contextual LFADS typically yielded a single fixed point per trial, facilitating interpretation of the inferred dynamics. However, this outcome is not guaranteed by the model. In principle, the network could learn more complex dynamics even in the presence of a per-trial bias. What we expect—and what we observed—is that introducing a contextual bias reduces the burden on the initial condition and time-varying input to encode condition-specific structure. Instead, the bias allows the model to parameterize a family of generators, often simplifying the resulting dynamics.

One limitation of our approach is that it assumes segments of neural activity with relatively stationary dynamics. In more complex tasks, modeling the per-trial bias as a piecewise-constant input could allow the generator to adapt to dynamics across task phases. Another challenge lies in hyperparameter selection—particularly in choosing the dimensionality of the bias—since decoding performance is often unaffected by the bias and interpretability is difficult to quantify.

## Acknowledgements

We would like to thank T5, T12 and their care partners. We would like to thank Beverly Davis, Kathy Tsou, and Sandrin Kosasih for administrative support. Support provided by Office of Research and Development, Rehabilitation R&D Service, Department of Veterans Affairs (N9228C, N2864C, A2295R, B6453R, P1155R, LRH); NIH-NIDCD (LRH), R01DC009899 (LRH), NIH-NINDS UH2NS095548 (LRH), NIH-NIDCD U01DC17844 (LRH), American Heart Association (LRH), NIH-NIDCD R01DC014034 (JMH), NIH-NINDS U01NS098968 (JMH), Wu Tsai Neurosciences Institute at Stanford (JMH), Larry and Pamela Garlick (JMH), Milton Safenowitz ALS Fellowship (NPS).

## Disclosures

The MGH Translational Research Center has a clinical research support agreement (CRSA) with Ability Neuro, Axoft, Neuralink, Neurobionics, Paradromics, Precision Neuro, Synchron, and Reach Neuro, for which LRH provides consultative input. LRH is a non-compensated member of the Board of Directors of a nonprofit assistive communication device technology foundation (Speak Your Mind Foundation). Mass General Brigham (MGB) is convening the Implantable Brain-Computer Interface Collaborative Community (iBCI-CC); charitable gift agreements to MGB, including those received to date from Paradromics, Synchron, Precision Neuro, Neuralink, and Blackrock Neurotech, support the iBCI-CC, for which LRH provides effort.

JMH is a consultant for Neuralink and Paradromics, is a shareholder in Maplight Therapeutics and Enspire DBS, and is a co-founder and shareholder in Re-EmergeDBS. He is also an inventor on intellectual property licensed by Stanford University to Blackrock Neurotech and Neuralink.

The content is solely the responsibility of the authors and does not necessarily represent the official views of the National Institutes of Health, or the Department of Veterans Affairs, or the United States Government.

CAUTION: Investigational Device. Limited by Federal Law to Investigational Use

## 6 APPENDIX

### 6.1 Variational Training Objective

The model is trained to maximize the evidence lower bound (ELBO) on the log-likelihood, summed over all data samples. The ELBO of the observed data sample *i* is given by:

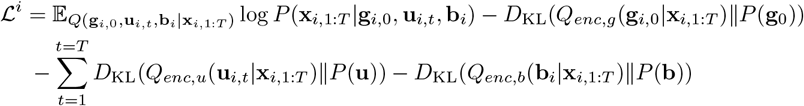

where the expectation is using the factorized distribution *Q*(**g**_*i*,0_, **u**_*i,t*_, **b**_*i*_| **x**_*i*,1:*T*_) = *Q*_*enc,g*_(**g**_*i*,0_|**x**_*i*,1:*T*_)*Q*_*enc,b*_(**b**_*i*_ | **x**_*i*,1:*T*_)Π_*t*_*Q*_*enc,u*_(**u**_*i,t*_ |**x**_*i*,1:*T*_). We assume isotropic Gaussian priors *P* (**g**_0_), *P* (**u**), *P* (**b**). Posterior distributions are parametrized by encoder networks. The data likelihood term corresponds to the Poisson log-likelihood of observed spike counts under generated rates.

### 6.2 Implementation and Training

Both the standard and modified LFADS models are implemented in Tensorflow 2 using GRU cells for the generator network. We use bidirectional RNN encoders for **g**_0_, **u**_*i,t*_, **b**_*i*_, and KL annealing during training. The dimensionality of the per-trial bias vector **u**_*i*_ is typically constrained to 2–10 dimensions.

Model optimization uses Adam with gradient clipping and early stopping. Monkey maze data was binned at 10 ms resolution and human iBCI data was binned at 15ms resolution. We regularize the generator dynamics via L2 weight penalties and restrict **u**_*i*_ through KL divergence and dimensionality limits.

### 6.3 Fixed Point and Linear Dynamics Analysis

To interpret the dynamical system learned by the RNN generator, we identify fixed points or slow points and analyze the linearized dynamics around them [17, 18]. A fixed point **g**\* of the generator dynamics satisfies:

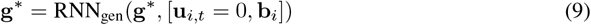

Since RNNs do not always yield exact algebraic solutions for fixed points, we identify *slow points* by minimizing the norm of the velocity vector:

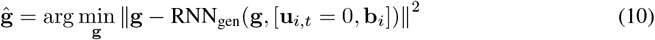

Optimization is performed using gradient descent from various initializations, seeded from latent states during real trials. Points for which the update norm falls below a fixed threshold (e.g., 10^*−*7^) are considered valid slow points.

To study local behavior near a slow point, we compute the Jacobian of the update function:

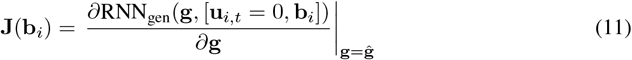

The eigenvalues and eigenvectors of **J**(**b**_*i*_) characterize the linearized dynamics:

- The real part of each eigenvalue indicates the stability of the corresponding mode (negative implies damping).
- Complex conjugate pairs indicate oscillatory modes, with imaginary parts determining the frequency and real parts controlling the rate of growth or decay.
- Eigenvectors (or generalized eigenspaces) define the oscillation plane or direction.

This analysis reveals how different trial conditions (via **b**_*i*_) induce shifts in fixed point locations, the orientation of eigenplanes, and oscillation frequencies, as shown in the pendulum and monkey reaching data.

### 6.4 Hyperparameters

All hyperparameters were set by optimizing ELBO on validation partition or manual quasi-optimization. The hyperparameters chosen for different analyses are listed in Table 3.

**Table 3:** Hyperparameters used to fit the contextual LFADS model for each dataset.

| Hyperparameter | Simulated Pendulum | Macaque Maze dataset | Human iBCI finger movement | Human iBCI Speech dataset |
| --- | --- | --- | --- | --- |
| Initial condition(I.C.) dimension | 5 | 128 | 10 | 128 |
| I.C. encoder hidden dimension | 128 | 128 | 128 | 128 |
| Time-varying (T.V.) input dimension | N/A | N/A | 1 | N/A |
| T.V. encoder hidden dimension | N/A | N/A | 128 | N/A |
| T.V. Input smoothness $\alpha$ | N/A | N/A | 0 | N/A |
| Bias dimension | 5 | 10 | 10 | 10 |
| Bias encoder hidden dimension | 128 | 128 | 128 | 128 |
| Generator hidden dimension | 200 | 200 | 200 | 128 |
| Dropout rate | 0 | 0 | 0 | 0 |
| Factor dimensions | 40 | 40 | 40 | 40 |
| L2 loss on trainable parameters | 0 | 0 | 1 | 0.1 |

### 6.5 Data augmentation for addressing non-stationarities in neural recordings

We outline the details of the non-stationarity correction method outlined in Section 4.3. The neural data consists of 966 trials with 52-54 trials for each of the 16 movements, and 107 trials without a movement.

Movements were cued with a continuous video of finger animation. For each trial, markers showed the final position of each finger and the fingers slowly moved from initial position to the target positions.

Cued target finger positions consist of two degrees of freedom (thumb vs all other fingers constrained to move together), and each finger group was cued independently to be flexed/extended/remain at rest. Trials alternated between fingers going from resting position to a target position and back. The iBCI participant was instructed to attempt to move his own fingers to follow the movement of the fingers in the animation. Multi-unit threshold crossings were recorded concurrent to attempted movements.

A linear finger-position decoder takes threshold crossings (**x**_*t*_) and predicts continuous finger positions (**p**_*t*_) at the corresponding time point: 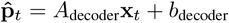. Our goal is to learn the decoder for the last quarter of the trials (trial numbers 724 *−* 966).

To evaluate different methods, data was randomly split into training (60% of trials), validation (20% of trials) and testing (rest of the 20% of trials). Note that trials in the target duration (trials 724 − 966) were present in both training and testing partitions.

Below we detail the steps for using contextual LFADS for data augmentation, by correcting the bias for each trial.

1. Identify the per-trial bias by applying the bias-encoder **b**_*i*_.
2. Divide the training trials into two sets based on trial ID, the non-target set 𝒮_1_ = [0 : 724] and the target set 𝒮_2_ = [724 : 966].
3. Compute the average bias for each set: 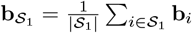 and 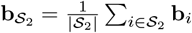
4. Apply bias correction for each trial *i* in the non-target set 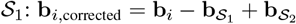. No change applied to trials in target set 𝒮_2_.

We applied identical steps for correcting the initial condition (yielding **g**_0,*i*,corrected_) for trials in the non-target set 𝒮 _1_.

The drift corrected neural activity was generated starting from the corrected initial condition as follows:

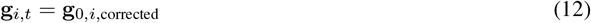

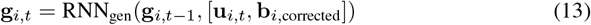

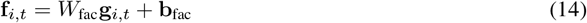

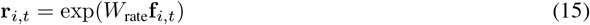

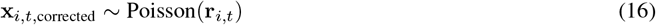

Finally, the corrected firing rate samples for trials in 𝒮 _1_ were combined with uncorrected firing rate samples for trials in 𝒮 _2_ to estimate the finger-position decoder. The decoder was subsequently evaluated on LFADS-smoothened firing rate on the test data.

